# Modeling the IL-2-Teff-Treg system in Systemic Lupus Erythematosus patients: possible mechanism and treatment strategies

**DOI:** 10.1101/2020.05.14.095513

**Authors:** Xin Gao, Xiaolin Sun, Jing He, Fangting Li

## Abstract

Systemic lupus erythematosus (SLE) is a non-organ specific autoimmune disease, which the pathogenesis of development is still unclear. For revealing the underlying mechanism, we construct a mathematical model depicting the interactions among CD4^+^ regulatory T cells (Treg cells), CD4^+^ effect T cells (Teff cells) and IL-2 in SLE patients to simulate and reproduce the development of SLE. Through our analysis, one possible pathogenesis is that the activation point and deactivation point of Teff in SLE patients produced a forward shift compared to the normal Teff. According to our simulation, a therapeutic window existed in the treatment of lupus with exogenous IL-2, which means there is a strict limit to the dose of IL-2 in clinic. Finally, we study three different dosing strategies and reveal that specific dosing regimens can better exert the effects of IL-2 and relieve the possible side effect of high dose IL-2. It is possible to promote the study of autoimmune diseases through our efforts and assist the development of precision medical treatment.

## Introduction

Systemic lupus erythematosus (SLE) is a complex and chronic systemic autoimmune disease^[1]^. Current therapy of active SLE mainly depends on corticosteroids and immunosuppressants due to the heterogeneous nature of this disease and lack of specific treatments ^[1, 2]^. In the past decades, studies showed that homeostatic impairment of CD4^+^ helper T cell (Th cell) subsets was crucial for the progress of SLE, which was characterized by downregulation of regulatory T cell (Treg) functions and overactivation of pathogenic effector T cells such as Tfh and Th17 cells ^[3–5]^. It has been revealed that IL-2 signaling deficiency contributed to the T cell impairment in SLE ^[3, 4]^. Recently, in our proof of concept clinical trial and some case studies, low-dose IL-2 supplementation was shown to be efficient in active SLE treatment and selectively upregulated Treg ratio and functions and decreased Tfh and Th17 cells^[6, 7]^. Further phase I/II clinical trials are underway to provide more details on the efficacy and safety of IL-2 therapy^[8]^.

IL-2 is a cytokine predominantly produced by T cells. Also named T-cell growth factor, it is traditionally regarded as a promoter of T-cell-dependent immune responses; however, recent studies prove that IL-2 is also essential to maintain functional Treg and thus control immune homeostasis^[9]^. These results have led to a series of studies all showing low dose IL-2 supplementation as a promising therapeutic strategy for a variety autoimmune-related diseases^[6, 7, 10, 11]^. All the low dose IL-2 application to date were reported to be safe and well tolerated, however, the minimum and maximum dose of IL-2 to stimulate Treg cells without activating effector T cells have not yet been established. Due to the poor safety record of high dose IL-2 therapy in cancer, it will be necessary to define the dose range and time course for low dose IL-2 administration to make the clinical application of low dose IL-2 efficient and safe.

The complex dynamic regulatory network of IL-2-Teff-Treg system in autoimmune disease makes it difficult to define the optimal dose and timing for low dose IL-2 therapy on autoimmunity in clinical studies or biological experiments. Mathematical modelling may facilitate precise quantitative analysis on this problem and direct future clinical or experimental studies. We established the mathematical model for IL-2-Treg-Teff regulatory system in SLE. We proposed three important results through this model. Firstly, this model exactly shows a therapeutic window and a bi-stable zone in most instances. With low dose IL-2 application, the proportion of Teff cells and the immune system tend to be homeostatic, but high dose IL-2 will saturate the IL-2 receptor of Tregs and activate Teffs. Subsequently, we construct and compare the healthy and SLE model to simulate the development of SLE. The forward movement of bifurcation point facilitate the generation of immune response with less antigen than normal case, which illustrates an important mechanism in pathogenesis. Lastly, The model demonstrated that different treatment strategies would affect the activation boundary of Teffs and the width of the therapeutic window. The IL-2-Treg-Teff modeling may help us better understand the pathogenesis and regulation of autoimmune diseases like SLE and provide certain guidance for clinical treatment.

### Overview of the IL-2-Teff-Treg system in the SLE patients

There are several feedback loops in IL-2-Teff-Treg system, which determines the development and pathogenesis of SLE (Fig. 1). As we all known, IL-2 has a decisive effect on immune system, immunity and tolerance by inducing the proliferation of Teffs. It proposed that interleukin-2 the T cells growth factor mostly secreted by Teffs and few by Tregs. In our IL-2-Teff-Treg system, Teffs release IL-2 while soluble IL-2 binds to IL-2R on the surface of Teffs and Tregs to promote the proliferation, despite most Tregs in patients are dysfunctional Tregs (named Tregs^dys^). IL-2 could also cause Tregs^dys^ transform to functional Tregs (named Tregs^func^), which inhibits and suppresses proliferation and activity of Teffs. In addition to cell-cell interactions, this system also contains the interactions of various intracellular cytokines. The self-antigens bind to T cell receptors after presenting by APC cells (antigen presenting cells) and phosphorylate NFAT (nuclear factor of activated T cells) to activate downstream pathways. One positive feedback loop in Teffs begins with IL-2 binding to IL-2R conforming IL-2/IL-2R complex then phosphorylating STAT5 (signal transducer and activator of transcription 5), which mediating the expression of IL-2R. Phosphorylated STAT5, however, also promotes the expression of Blimp1 and thus inhibits the release of IL-2. Except Tregs produces few IL-2 that we neglect Blimp1 in Tregs, the basic pathways were similar to those in Teffs. Besides, our model differentiates Tregs^dys^ and Tregs^func^ directly according to the concentration of phosphorylating STAT5 and also include the competition effect of soluble alpha chain of IL-2R (sCD25).

**Fig. 1.**
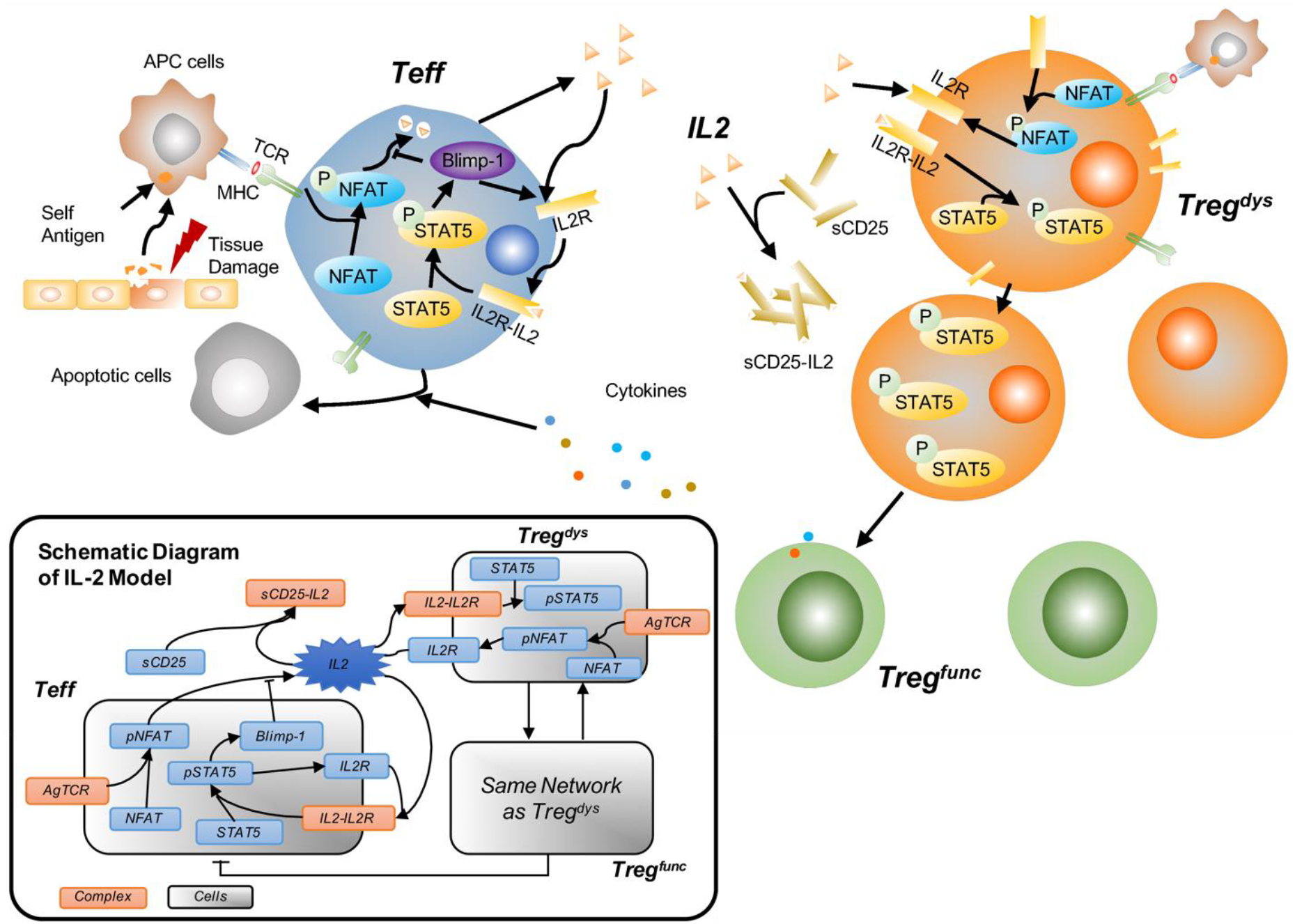
The molecular mechanism of IL-2-Teff-Treg system. It demonstrates the interaction of various cells and cytokines in IL-2-Teff-Treg system. The black arrows represent activation or facilitation, when the lines end by horizontal lines indicate inhibition or suppression. In the presence of sufficient antigens in vivo, IL-2 binds to the receptor on the surface of T cells to promote proliferation and enhance downstream response. The IL-2 pathway is the same in both Teff and Treg cells. The only difference is that Tregs secrete little IL-2 compared to Teffs, which can be ignored in simulation. Two main sources of IL-2 including autocrine and exogenous supply existed in our model, but there is still a certain amount of IL-2 background concentration. It needs to be emphasized that although the number of Treg is only one tenth of Teffs, the density of surface receptors is about a hundred times, in general, Tregs are more competitive than Teffs. The schematic network diagram of IL-2 system is depict in the lower left corner.

Recent progresses show that the Treg-Teff system in SLE patients have functional defects^[12]^, which causes great differences of Treg cells between patients and normal people in activity ^[7, 13]^. The fact most of the Treg cells in the SLE patients are in a low active state ^[7]^ suggests that the Treg cells in SLE patient disable to regulate Teffs activity when Teffs overactivated. Apart from this, several work showed that concentration of IL-2 in peripheral blood of SLE patients is lower than normal level^[14]^. We also tested the number of Treg, Teff, CD8^+^, DNT, NK and NKT cells in peripheral blood of SLE patients as well as the healthy controls (Fig. 2A). The IL-2 producing immune cells of patients counts about half the number of healthy people, indicating that the IL-2 concentration in patients is probably lower than the control. Unfortunately, due to the limitation of the instrument measurement accuracy, the IL-2 concentration is difficult to measure directly. Taken together, it is reasonable for us to assume that the development of SLE as well as the treatment on SLE patients with IL-2 (other autoimmune disease by extension) are probably related to the transformation of Treg^dys^ to Treg^func [15]^.

**Fig. 2.**
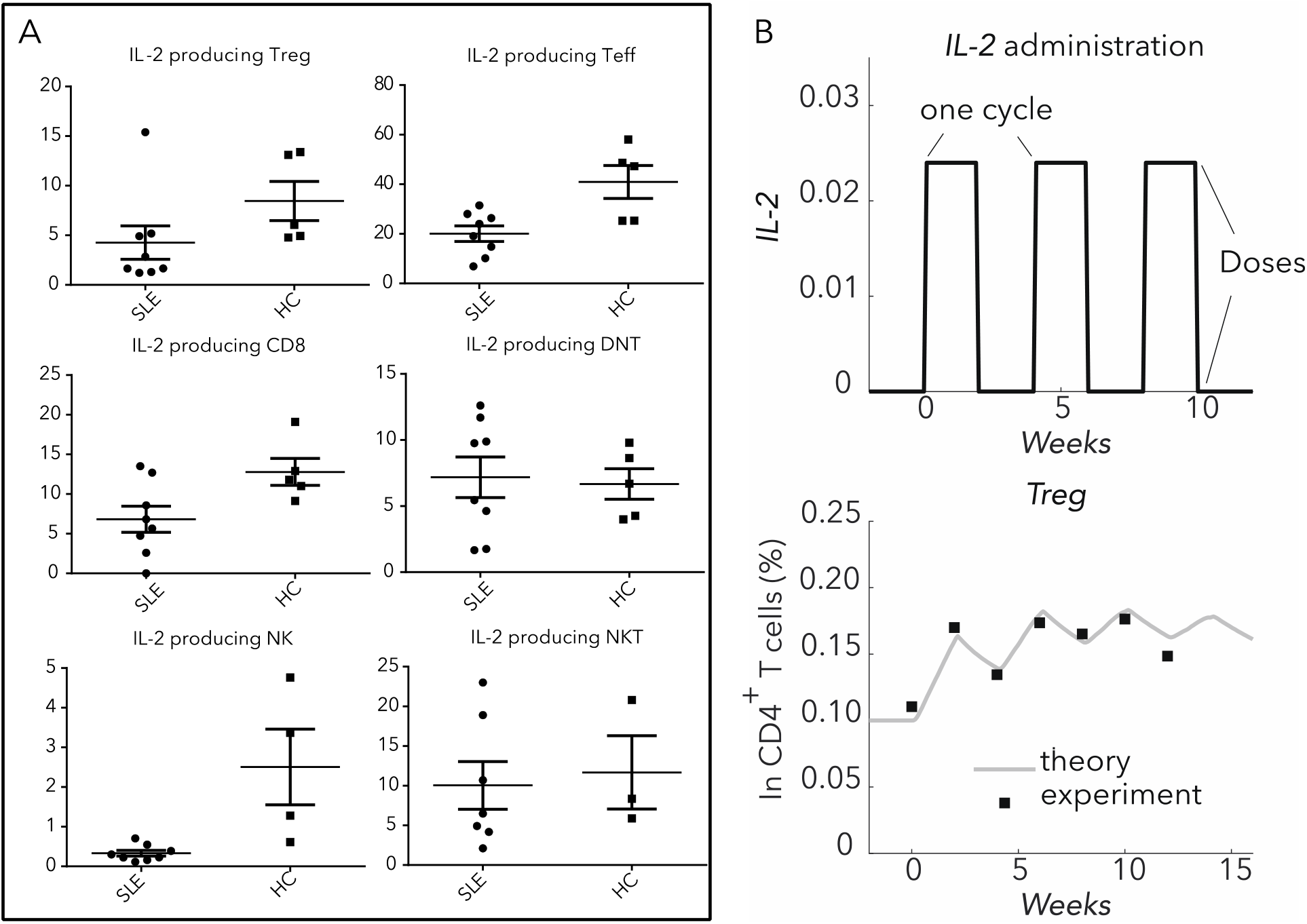
Comparison of various immune cells between SLE patient and healthy control. (A) Counts of IL-2 producing immune cells including Treg, Teff, CD8^+^, DNT, NK and NKT cells are shown for SLE patients (8 samples, n=8) and the healthy control (5 samples, n=5). Relative proportions of IL-2 producing CD8^+^ cells and Teff subsets in peripheral blood were analyzed by flow cytometry. Each single black dot indicates the percentage of the corresponding cell subset among immune cells of one single SLE patients or healthy control. Horizontal lines in scatter plots indicate the median and interquartile range. (B) The results of our theory matches well with experimental data ^[2]^ under the periodic IL-2 administration. Upper panel demonstrates the pattern of periodic IL-2 administration while the bottom panel shows the fitting result of the proportion of Treg cells in CD_4_^+^ T cells. The gray line represents our simulation, while the square black dots indicate the experimental data.

### Mathematical model of IL-2-Teff-Treg system in SLE patients

Based on the data analysis and literature research, we established a model involving the interaction of various types of T cells and other intermolecular interactions. We supposed that the majority of patients’ Treg cells are inactive or dysfunctional regulatory T cells (Treg^dys^) and could not play the regulatory role in SLE system unless they transform to active state. The active or functional regulatory T cells (Treg^func^), however, may also randomly transit to the inactive state. For the sake of simplicity, we set the rate of this reverse transition to a constant value in simulation. There are mainly two processes that affect the proliferation of Teffs. The first is competition, where other cells or cytokines with IL-2 receptors can reduce the IL-2 obtained by Teffs through competition. For examples, soluble CD25 binds IL-2 to form a complex, and simultaneously the IL-2 receptor on the surface of Tregs also compete to bind IL-2 to activate the downstream pathways. In addition to the competition, by producing cytokines like IL-10, IL-35, and TGF-β, Tregs can directly suppress the activation of Teffs. Other pathways like releasing granzyme B and perforin 1 and combining with APC cells to induce tryptophan can also reduce the numbers of Teff cells. Briefly, Tregs can effectively regulate Teffs through direct inhibition and competition.

To highlight the key points, we simplify our mathematical model from the following aspects. Firstly, by considering the number of Teff cells is much more than other immune cells subsets^[16]^, we assumed that the endogenous IL-2 is all secreted by Teff cells. Secondly, we approximate each edge (Edge represents interaction between two nodes connected by this edge.) in Fig. 1 an elementary reaction. For example, it is displayed in our model that Teffs proliferation will directly lead to the up-regulation of IL-2, yet actually several kinds of molecular might exist during this pathway, such as X-chromosome–defined signaling lymphocytic activation molecule-associated protein (SAP) and calcium (Ca^2+^)^[17]^. These kinds of molecular are indeed important, however, during our model, they mainly work as the intermediary. Hence to avoid a bloat model, they have all been omitted. Thirdly, Teff cells in our model is a collective term for all effector T cells which actually could be divided to follicular helper T cells (TFH), Th17 cells and etc. IL-2 can via STAT5 constrains Th17 cells^[18]^, it can also limit TFH differentiation by up-regulating Blimp-1 as well as down-regulating Bcl-6^[19]^. To put it into a nutshell, all of these pathways are explaining how IL-2 down-regulates T effective cells in detail, and the direction won’t be changed if these details are omitted. Therefore, we ignore these details in our model and suppose it is reasonable to consider the Teff cells as a whole.

Our ordinary differential equations consist of 15 variables: IL-2R, IL-2/IL-2R, pNFAT, pSTAT5 in Treg and Teff respectively and Blimp1, IL-2, σ_h_, σ_r_, σ_r_^a^, sCD25, sCD25IL-2. The detailed meanings of these variables given in the table below. These equations are:

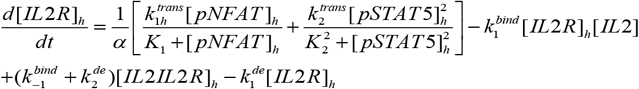

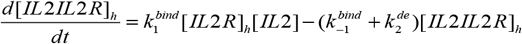

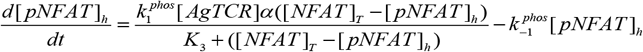

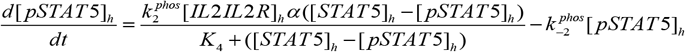

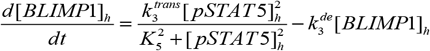

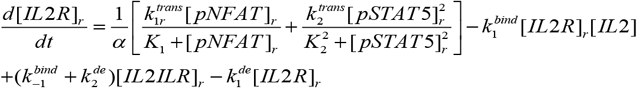

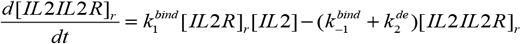

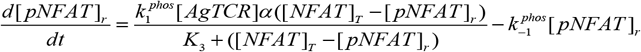

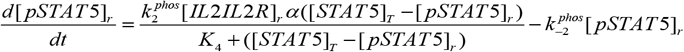

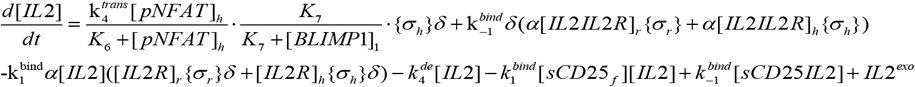

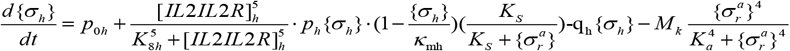

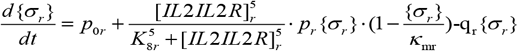

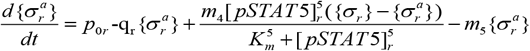

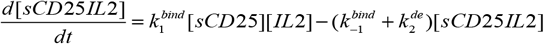

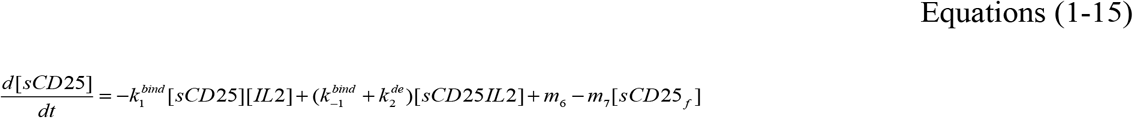

The first two variables represent the areal density of IL-2 receptor and IL-2/IL-2R complex. Considering IL-2 receptor pathway was regulated by STAT5, we add a conversion factor α in the front of its generate item, and the negative item in the first two equations represent degradation and dissociation. Equation 3-5 represent three intracellular cytokines and each equation consists of phosphorylation item and degradation items. It should be noted that AgTCR is the surface antigen receptor complex and is saturated in most cases. Equation 6-9 have same meaning as Equation 1-4 except in Treg cells. Equation 10 represent concentration of IL-2 in full space, which can be produced by Teffs or injected. Equation 11-13 represent number density of Teff cells, active Treg cells and total Treg cells respectively, that is, all these cells will proliferate under the action of IL-2 and apoptosis at a certain rate. The last two items characterize the competitive effect of free IL-2 receptors on IL-2 which is easy to understand. The label ‘[…]’ denote concentration or areal density of variables and ‘{…}’ denote number density of cells. All these variables are positive and the unit, meanings and initial values of these variables are list in table 1.^[20]^ Thanks to the gorgeous work by hofer et.al^[21]^, parts of parameters are relatively reliable from experiment. Other parameters are given based on fitting and estimation form medical analysis. The purpose of this complex model is to simulate as many possible scenarios as possible in immune system by considering most details we known.

**Table. 1.**
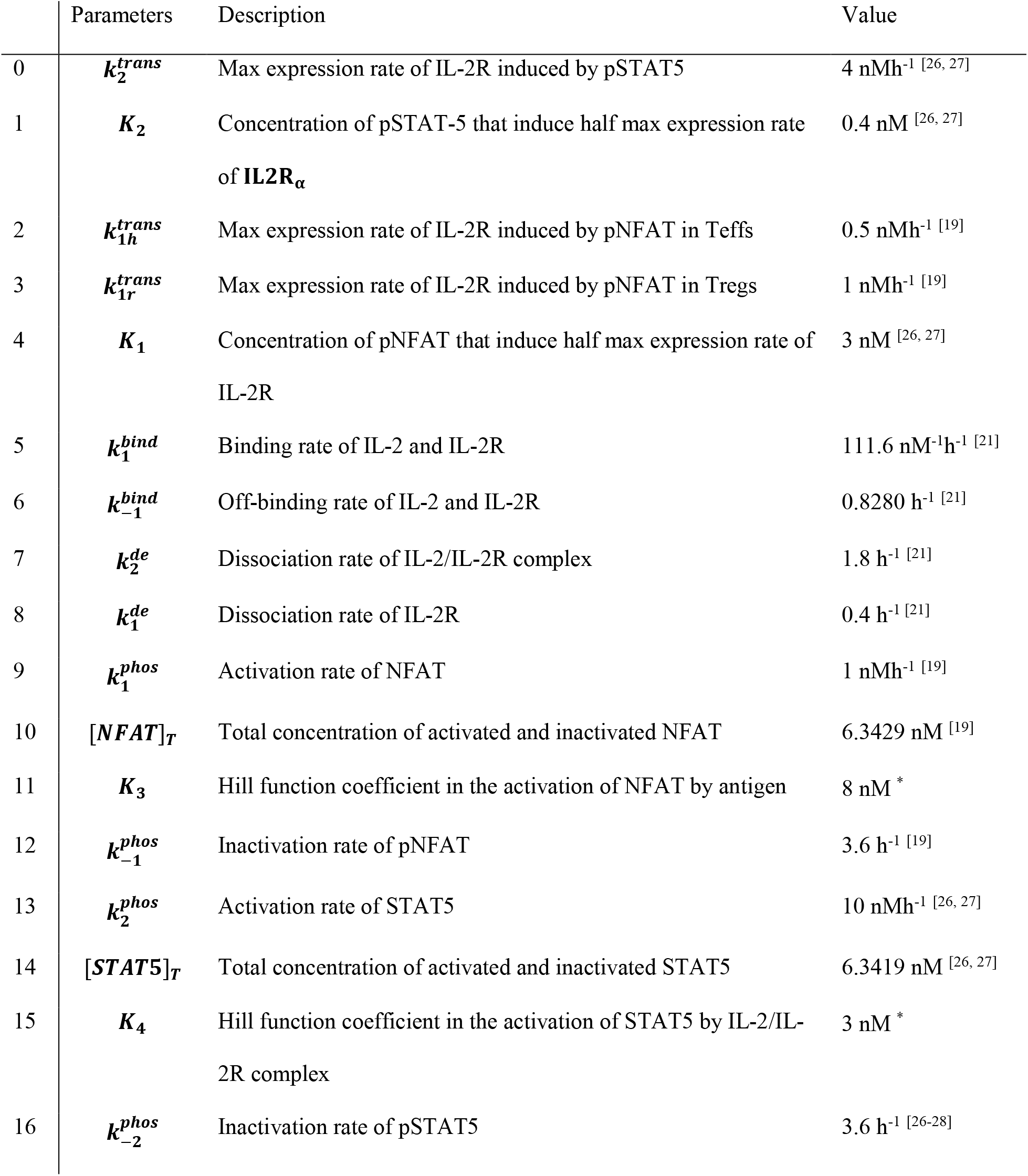
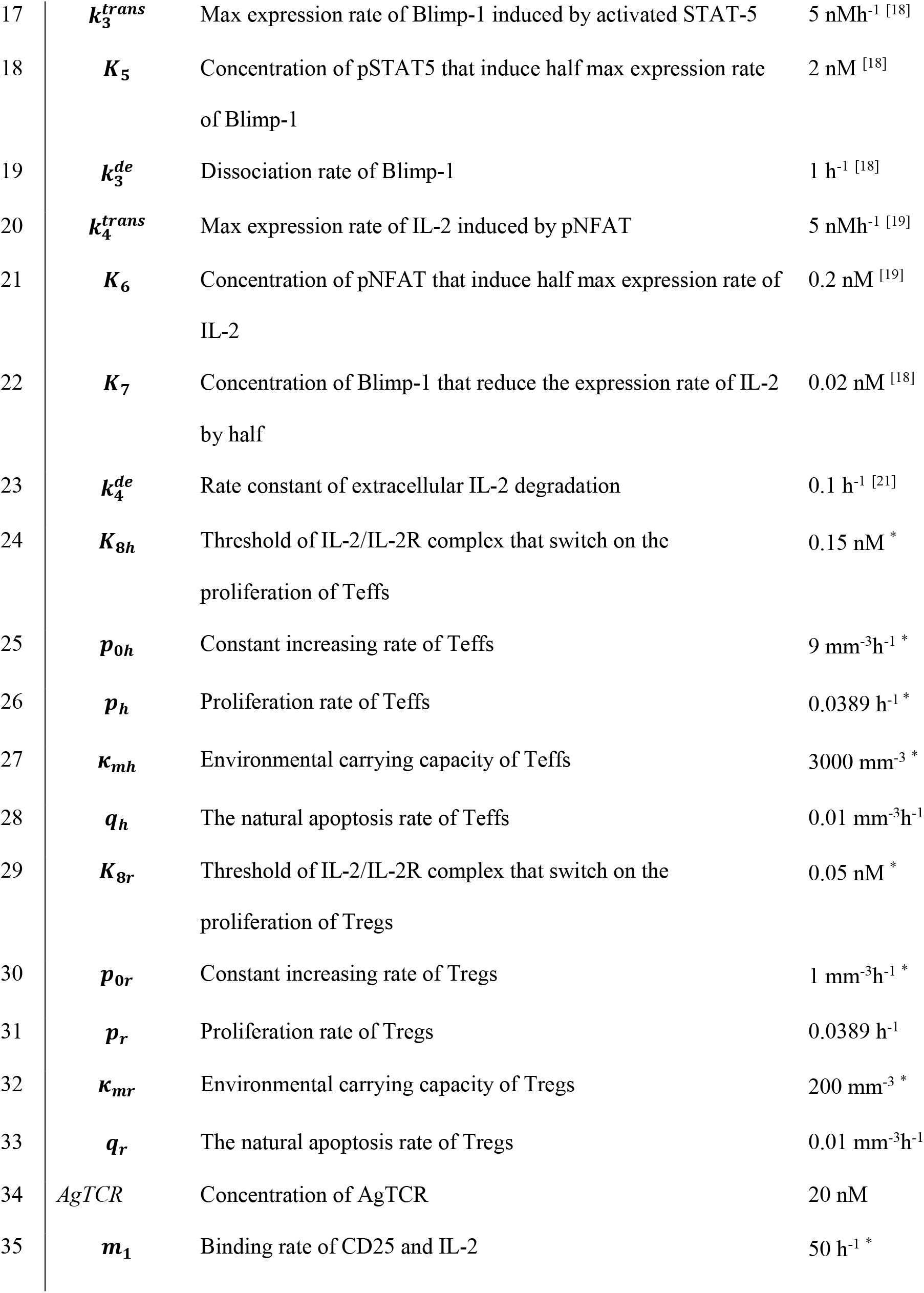
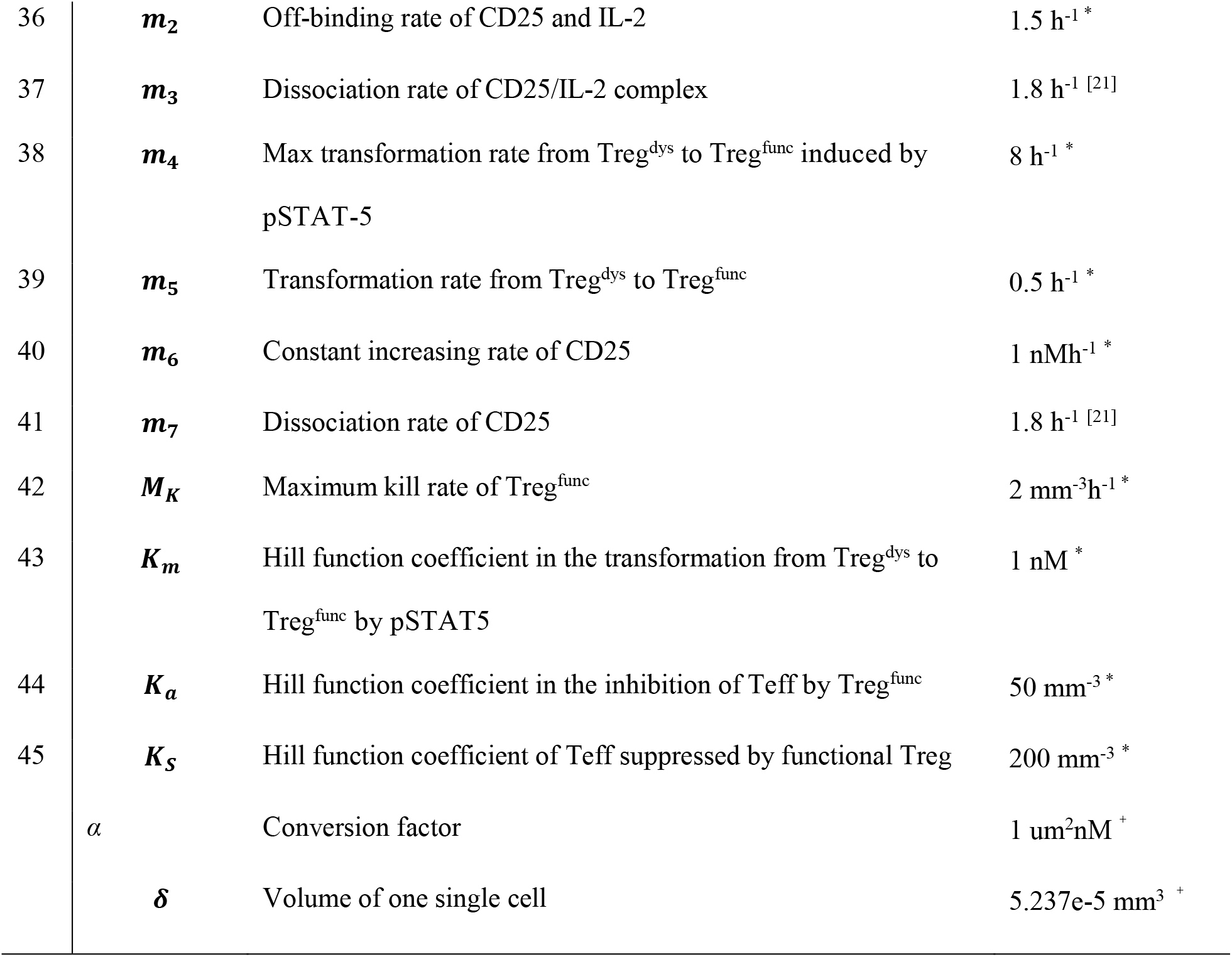
List of All Parameters A set of standard parameters are list in table. 1. Part of these parameters are introduced from other work on immunization after a minor modification when the other come from our estimation or calculation. The label ‘*’ represents that the corresponding parameter is estimated based on actual conditions. The label ‘+’ represents that we calculate this value based on our model or actual situations.

We simulated the kinetic response of immune system in the blood and this system has a definite volume V_T_ (The volume of T cells couldn’t be ignored in lymph nodes compared to the blood). There are two types of cells in this system, Tregs and Teffs, which Treg cells have two status: functional and dysfunctional. In a resting environment, IL-2 secreted by Teffs will fall out of the gradient. However, in the flowing blood, we can think that IL-2 secreted by Teffs is homogeneous in the system. The variable σ_h_ represents the number density of Teff cells as σ_r_ represents the number density of Treg cells, such volume of all Teff cells can be written by σ_h_V_T_V_S_ and volume of all Treg cells is σ_r_V_T_V_S_, which Vs denotes the volume of one single cell. We assume that all the cells are evenly distributed in system and have a large spacing compared to the volume of cell so that the probability that each IL-2R will collide and bind to IL-2 will be approximately same no matter in the surface of Treg or Teff (Fig. 1).

For complex biological systems, several assumptions were introduced to make the model easier to understand and handle. Firstly, we assume that the patient’s Tregs only have functional and dysfunctional states, although there may be some middle status between completely functional and dysfunctional. Secondly, we assume that the volume of single Treg cells (VS) approximately equals to the volume of single Teff cells. Then, although Treg could secrete IL-2 in few instance, it is negligible in the patient’s blood. In addition, we use celebrated Hill function in most of equations to simulate the instance that the effect of one variable on another variable will saturated. Because of the difference in the units between different variables of the equation, we divide the equation into three parts to derive and interpret.

The first is the interactions of receptors on surface and intracellular factors. Before we derivate the equations, it is convenient to derive the conversion factors for the interaction of receptors on the surface with intracellular cytokines first. As we described above, IL-2 receptor and IL-2 complex can activate intracellular pathways and phosphorylate STAT5, such interactions can be simplified if we assume these surface receptors are evenly distributed intracellular. The unit of areal density of surface receptors is written as um^−2^ and the concentration of intracellular cytokines is written to be nM. (The volume of single T cells is approximately 1000um^3^ and the radium is nearly 10um) The conversion factor is simply satisfied:

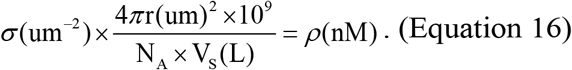

We defined 4 as α. So the formula above can be simplified as

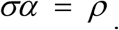

For example, the item describe the production of IL-2R in Teffs can be written as:

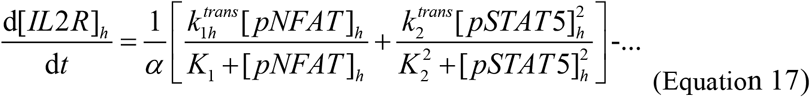

This item describes the generation of IL-2 receptors on surface, which [pNFAT] in Teffs have different units with [IL-2R], so that we divide generation item by α to simulate the process that transport IL-2R to the surface after generating. Similarly, the item describe the activation of downstream pathways of receptor complex on surface should multiple by α to make sure they share the same dimension with intracellular cytokines.

The second class is the item of IL-2 secretion. It is clear that the IL-2 added to the system is all secreted by Teffs during a very short time dt,

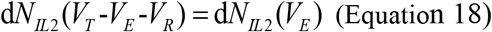

where d*N*_*IL2*_ (*V*_*E*_) is the incremental IL-2 produced by Teffs and d*N*_*IL2*_(*V*_*T*_−*V*_*E*_−*V*_*R*_)is the incremental number of extracellular IL-2. We are concerned with the concentration of [IL-2] in whole system, so we divide both sides of the equation by (V_T_−V_E_−V_R_). Therefore, the concentration of [IL − 2] is given by

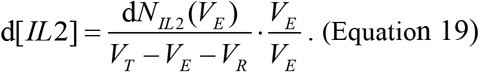

Obviously, d*N*_*IL2*_ (*V*_*E*_)/*V*_*E*_ represents the incremental concentration of IL-2 produced by Teffs and it can be replaced by the generation item:

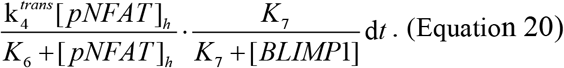

It should be noted that *K*_7_/(*K*_7_ + *Blimp*1) is corresponding to the inhibition of Blimp-1. Using

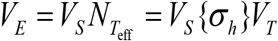

to do a simple replacement and using σ_*h*_ to represent the number density of Teffs in order to simplify the expression of the equation. [IL-2] is given by

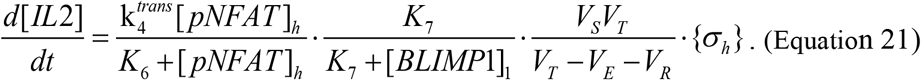

We simplify *V*_*S*_*V*_*T*_/(*V*_*T*_ − *V*_*E*_ − *V*_*R*_) to δ in our model and can derive that the volume-related factor δ existed on every terms involving interactions between cells and intracellular factors. The last class items consist the collision of IL-2 with surface IL-2 receptors. As we derived before, there are totally N cells distributed evenly in the system that VT indicates the whole volume of our system and Vs indicates the volume of single cell, which R is the radius of the cell. We consider the probability of a single IL-2 molecule colliding with a single cell and here we view IL-2 approximately at a particle point. The free space belonging to one single cell can be written as

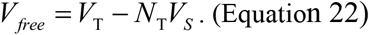

In the stationary reference frame relative to IL-2, each single cell moves at a relative speed ***v***_***ri***_ = ***v***_***i***_ − ***v***_***IL***−***2***_. In interval dt, the volume covered by the movement of cell i is

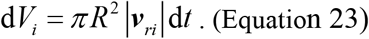

IL-2 molecule can appear at any place in free space with same probability. When IL-2 overlaps with the space dV_i_ covered by T cell in dt time, we can view that IL-2 collide with this cell. The probability that IL-2 collide with the cell in dt time is written as:

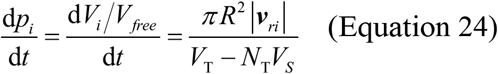

It should be emphasized that we default the cell did not collide with other cells in dt time. In fact, We can always find the interval that adjacent cells will not collide with each other within a limited time as long as V_free_>0. Then the total number of n IL-2 molecules colliding with one single cell in dt time is

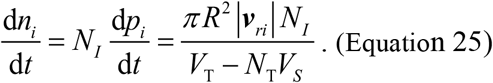

Notice that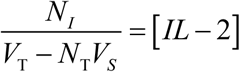, we can simplify the equation above and get the average times of collision.

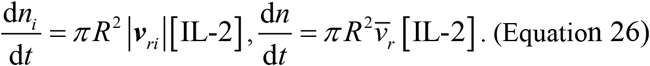

We have derived the number of collisions between single cell and IL-2 molecules per unit time. Let us denote the reaction cross section of IL-2 and IL-2R to b and the total number of IL-2R on the surface to N_IR_. Similarly, IL-2 molecules will appear at any place on the surface with same probability that the probability of reaction is

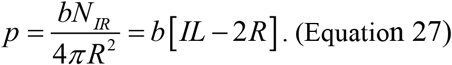

Finally, the times that IL-2 collides with IL-2R per unit time is

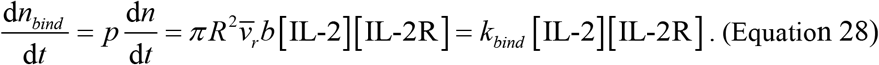

## RESULT

Applying our model to some specific immune diseases is one of our goals to construct this immune response model. For example, He et al. (2016) use interleukin-2 therapy to treat systemic lupus erythematosus (SLE). As a non-organ-specific autoimmune disease (AID), however, there is currently no perfect therapy with no side effect. For improving our understanding of specific mechanism of SLE and better developing treatment options, we constructed this SLE model to characterize the role of IL-2 in system and proposed a therapeutic window of IL-2 treatment. We also demonstrated that it will have a broader therapeutic window of IL-2 by adjusting some parameters and will be more effective by changing the mode of IL-2 administration.

To verify the reliability of our model, we introduced the data from He et.al 2016 and compared theoretical ratio of Tregs to the patients ^[2]^. They treated patients with autoimmune and inflammatory conditions by low dose rhIL-2 (10,000, 30,000 or 100,000 IU) daily for two week then stop for two weeks and detected the proportion of Tregs in CD4^+^ T cells every two weeks. The result of simulation demonstrated that the proportion of Tregs in CD4^+^ T cells increased in the form as wave within first ten weeks after treating by IL-2 and then decreased slightly. In Fig. 2B, it significantly shows that our theory matches satisfactory with the clinic data.

### Modeling the role of IL-2 in the treatment of the SLE patients: a double-edged sword

Based on our model, we analyze how the IL-2 interacts with Teff and Treg cells. We consider the proliferation of Teff and Treg cells, the increasing of IL-2 receptors on surface, the competition between Teff and Treg cells for IL-2, and the suppression and killing effect of Tregs. Because of the high density of IL-2 receptors on Tregs, once IL-2 released into the system, it will bind to the Tregs preferential, then phosphorylating STAT5 in Tregs to trigger the activation and proliferation of Treg cells. However, the low dose ensures most of IL-2 binding to the receptor on the surface of Tregs rather than Teffs due to its weak competition. Then Tregs transform to functional state, and suppress Teff cells through producing immunosuppressive factors like interleukin-10 (IL-10), transforming growth factor β (TGF-β) and other cytokines.

Unfortunately, when facing higher dose of IL-2, Teffs can also compete IL-2 and proliferate, which aggravates the condition of the illness. When IL-2 is excessive, receptors on the surfaces of Teffs and Tregs can bind IL-2 to form complex without competition and activate downstream response pathways to promote self-proliferation. Moreover, although dysfunctional Tregs can partly transform to functional Tregs, the suppression is not enough to counterweigh the rapidly self-proliferation of Teffs. According to this effect, we divide IL-2 dose into three classes by the corresponding ratio of Teffs to Tregs and the number density of Teffs (Table 3) by the fact that ultra-low has no obvious effects on the system, low dose IL-2 reducing the number of Teff cells when high dose increasing it. However, it is rather hard to control the dose precisely in clinical experiment because of the specificity among patients. Various kind of disasters may occur in clinic if we choose the wrong dose of IL-2, for example, ultra-low dose is not adequate to turn the switch on to activate dysfunctional Tregs, simultaneously high dose IL-2 will activate Tregs as well as Teffs and worsen the situation (Fig. 3A). Recent research shows that high dose IL-2 increases the rate and grade of graft versus host disease (GVHD) in allogeneic stem cell transplantation^[15]^. However, the mechanism of severe side effects with high dose IL-2 in autoimmune diseases has not been fully elucidated.

**Fig. 3.**
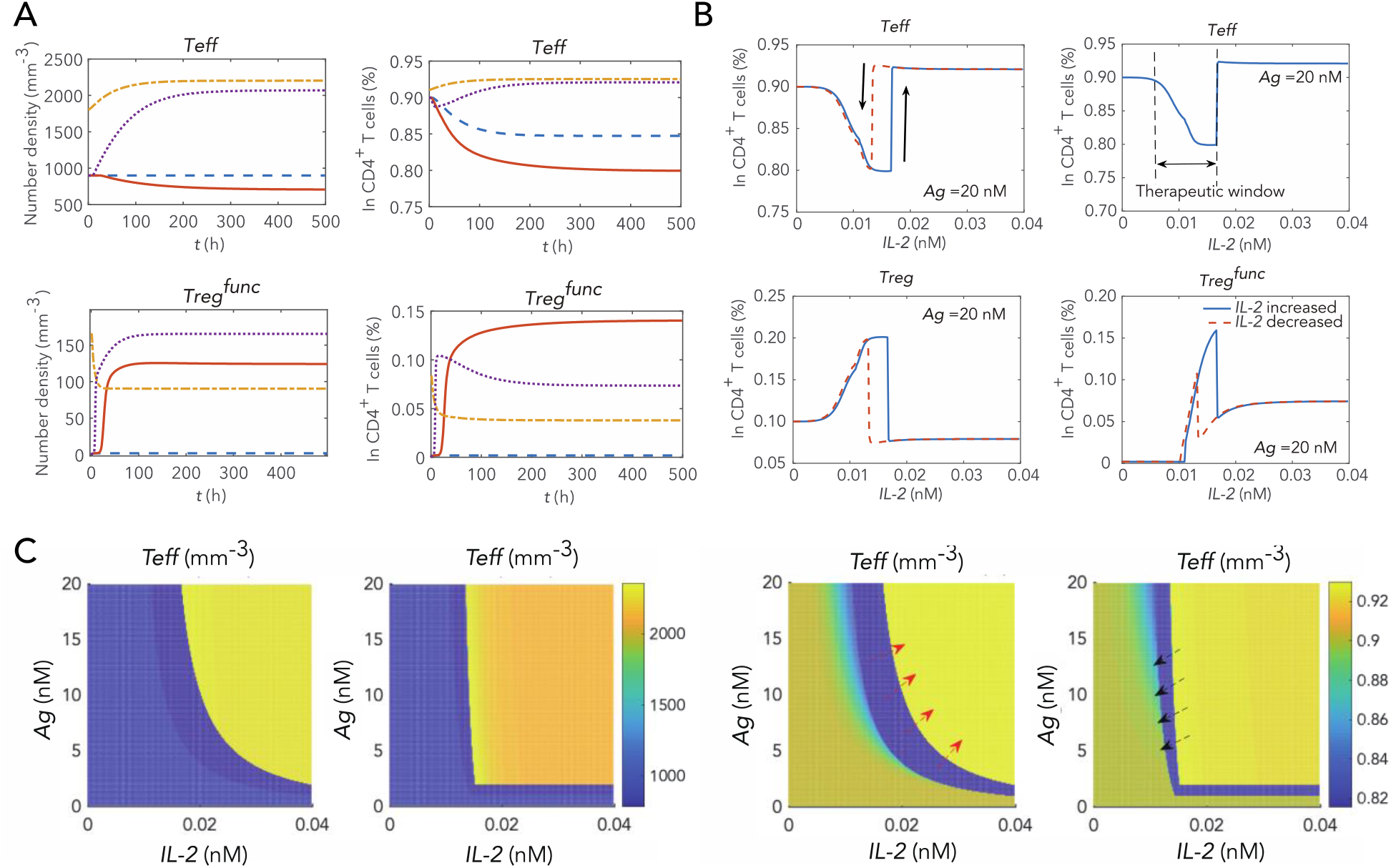
Different doses of IL-2 cause various immune responses as well as antigens in SLE treatment. (A) These images show the evolution of IL-2R, cell density of Teff and functional Treg under different concentrations of IL-2 (0.010 nM, 0.015 nM, 0.020 nM and 0.030 nM). Ultra-low dose IL-2 (0.010 nM) neither makes dysfunctional Treg transform to functional Treg nor proliferating Teff in large-scale. Low dose IL-2 (0.015 nM) in our model is located in therapeutic window meanwhile high dose IL-2 (0.020 and 0.030 nM) enables IL-2 receptors on Treg and Teff to bind sufficient IL-2. It is worth to mention that there is still another steady-state under low dose IL-2. (B) Ratio of Teff cells, Treg cells and functional Treg cells in CD_4_^+^ T cells, with increasing IL-2 in blue line and decreasing IL-2 in red dot line respectively. Teff has a bi-stable state in the attachment of the therapeutic window, which may narrow the original window under the external noise. (C) Cell density and ratio of Teffs vary with exogenous IL-2 and antigens. The left two pictures show ratio of Teff in CD_4_^+^ T cells with increasing and decreasing IL-2 respectively, while the right two charts show the changes in their cell density. We can intuitively observe the shift of therapeutic window with antigens and IL-2 from the set of figures. It indicated that there is a relatively wide therapeutic window when treating SLE patients with IL-2 and that it is difficult to fall back into the therapeutic window once a patient is treated with a high dose IL-2.

By modelling this double-edged sword effect of IL-2 by adding different dose of IL-2, we can easily judge from the results (Fig. 3B) that only the dose of IL-2 in a certain range can reducing the ratio of Teffs, which approximately equals to the range of the activation point of functional Tregs and the over-activation point of Teffs. As what we have deduced qualitatively, it causes the proliferation of almost all types of immune cells when IL-2 is excessive and that is why using high dose IL-2 to treat tumors. We also proposed that once using excessive dose of IL-2 in treatment, the therapeutic window will be narrower than normal case when ceasing suppling IL-2 (Red dot lines in Fig. 3B). This phenomenon reminds us the significance to control the amount of IL-2 accurately in clinic.

### The Development and Pathogenesis of SLE patients

In order to explore the development and pathogenesis of autoimmune diseases similar to SLE, we simulated the kinetics of patients and normal healthy person under the same dose of IL-2/Ag. We make a sight modification including parameters, initial state and equation to the original SLE model and construct a healthy model for comparison (Fig. 4A). First, because of the over-activated Teffs in SLE patients, it is rational to modify the value of max expression rate of IL-2R from 0.5 to 0.1. Second, considering all Tregs in healthy model are functional Tregs, we remove the impact caused by dysfunctional Tregs in our SLE model. Third, the initial value of variables in normal person is different from the patient. Under sufficient antigens/IL-2, normal person can also loss the healthy balance of activity between Tregs and Teffs, so it is reasonable to take the final state of healthy kinetic under sufficient IL-2/antigens as the initial state of SLE model. The detailed values of all variables are list in table2.

**Fig. 4.**
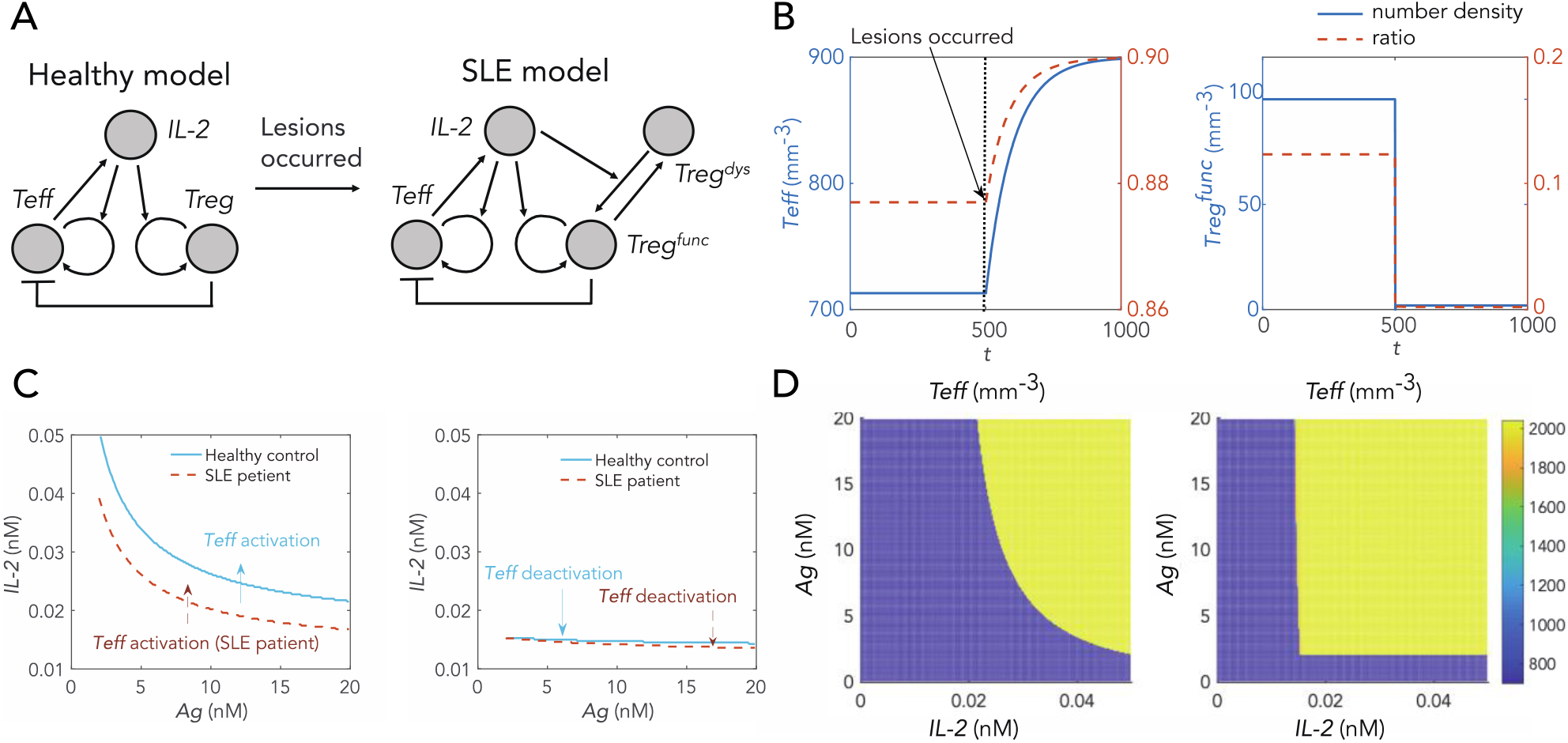
Pathogenesis of SLE compared with healthy model. (A) Schematic diagram of healthy control. It proposed that this toy model can clearly show the interactions between Teff and Treg cells. In original SLE model, we assume that Tregs in vivo are all dysfunctional in initial and Teffs have more competitive receptors. In the healthy model before the damage, however, all Tregs were functional Tregs. (B) We simulate the evolution of Teff and functional Treg number density over time after the structural lesions occurred. (C) Left panel demonstrates Teff activation boundary of healthy person and SLE patients respectively. Obviously, in SLE patients, Teff is activated with less antigens as well as IL-2 comparing to healthy person. Right panel indicates the Teff deactivation boundary of SLE patients and healthy person, which means Teff is little hard to deactivate than healthy control. (D) Proportion of Teffs in all CD4^+^ T cells vary with increasing (left) and decreasing (right) exogenous IL-2 in healthy person. The therapeutic window of healthy person, however, is obviously wider than that of SLE patients which shown in Fig. 3c. The activation and deactivation boundary of SLE patients and healthy control are compared in Fig. 4c.

**Table. 2.** Value of all variables in development and treatment of SLE with sufficient antigens

| | $[IL2R]_h(\mu\text{m}^{-2})$ | $[IL2IL2R]_h(\mu\text{m}^{-2})$ | $[pNFAT]_h(\text{nM})$ | $[pSTAT5]_h(\text{nM})$ | $[Blimp1]_h(\text{nM})$ |
| --- | --- | --- | --- | --- | --- |
| Initial value | 0 | 0 | 0 | 0 | 0 |
| Final value(HC) | 0.2973 | 0.0014 | 1.9649 | 0.0026 | 0 |
| Final value(SLE) | 0.4966 | 0.0029 | 1.9649 | 0.0055 | 3.8e-5 |
| | $[IL2R]_r(\mu\text{m}^{-2})$ | $[IL2IL2R]_r(\mu\text{m}^{-2})$ | $[pNFAT]_r(\text{nM})$ | $[pSTAT5]_r(\text{nM})$ | $[IL2](\text{nM})$ |
| Initial value | 0 | 0 | 0 | 0 | 0 |
| Final value(HC) | 0.9943 | 0.0047 | 1.9649 | 0.0088 | 0.0001 |
| Final value(SLE) | 0.9972 | 0.0059 | 1.9659 | 0.0111 | 0.00014 |
| | $\sigma_h(\text{mm}^{-3})$ | $\sigma_r(\text{mm}^{-3})$ | $\sigma_r^f(\text{mm}^{-3})$ | $[sCD25IL2](\text{nM})$ | $[sCD25](\text{nM})$ |
| Initial value | 700 | 100 | 100 | 0 | 0 |
| Final value(HC) | 713 | 100 | 100 | 0.0093 | 0.5556 |
| Final value(SLE) | 899 | 100 | 2 | 0.0011 | 0.5556 |

**Table. 3.** Therapeutic effect of different doses of interleukin-2 in our simulation.

| IL-2 dose | *Treg* activation | *Treg*<sup>func</sup> activation | *Teff* activation | Ratio of *Treg*<sup>func</sup> | Ratio of *Teff* |
| --- | --- | --- | --- | --- | --- |
| Ultra-low | Yes | No | No | ~0% | ~85% |
| Low | Yes | Yes | No | ~15% | ~80% |
| Medium | Yes | Yes | Yes | ~7% | ~92% |
| High | Yes | Yes | Yes | ~3% | ~92% |

For researching the development of systemic lupus erythematosus, we inflict structural damage on the system at a certain time and transform the healthy model to SLE model. According to Fig. 4B, Teffs show a significant increase in both number density and proportion after systemic injury, when functional Tregs show a significant decrease. Revealing the intrinsic discrepancy between SLE patients and normal person in immune response is the original intention of our modeling. For that reason, we take the same analysis method in SLE patient to normal person and depict the bifurcation graphs with changes in Ag/IL-2 parameters. Left panel in Fig. 4D show the trend of Teffs, Tregs, proportion of Teffs with increasing IL-2 respectively, when figures in right panel correspond to the circumstances with decreasing IL-2. Comparing the circumstances of normal person to SLE patients, it demonstrate that the Treg activation boundary and the Teff over-activation boundary of SLE patients move forward compared to normal person (Fig. 4C). This means that the patient’s immune tolerance is worse than that of a normal person, and once patients is over activated it is more difficult to reverse to a ‘healthy’ state than a normal one.

For a normal person, as IL-2/Ag increases, Tregs proliferate early than Teffs and release cytokines like IL10 to ensure that the body has a certain degree of immune tolerance. When IL-2 decreases, Teff first returns to the normal level and then Tregs. Teff, however, may be auto-proliferate in patients, before Treg when Treg has functional defects. In this section, we established two idealized models to simulate the kinetic responses of normal and patient under a stimulation of exogenous IL-2 or Ag. Through simulation, we found that SLE patients have worse tolerance than normal person and the sequence that Tregs increase first and then Teffs may be broken which causes the disorder in vivo.

### Broaden the therapeutic window of IL-2-Treg-Teff system

As we shown in Fig. 3B, we can got a satisfactory result by precisely choosing an appropriate dose of interleukin-2. However, due to the significant variation of physical and sexual characteristics among different patients, the appropriate dose of IL-2 of wide range of patients are not always the same. If we take the ultra-low dose IL-2 that may has little effect, but choosing an excessively high concentration IL-2 causes receptors on Treg surface saturated and makes Teffs over-activated. Simultaneously, without competition, Teffs will inevitably over-proliferate until the environment carrying capacity reaches the upper limitation. Individual traits like race, gender impact the dynamics and characteristic of patients. Specifically, in our SLE network, due to the prevalence of diversity among patients, part of our parameters and the initial value of variables will be different. In other words, investigating the influence of parameters on system dynamic properties like the width of therapeutic window is of great significance to the development of precision medicine.

Fig. 5A shows the therapeutic windows for individuals at three concentration of antigens treating with increasing dose of exogenous IL-2. Differences between these three therapeutic windows indicate immune system are not robust to part of parameters and it makes great sense to analyze sensitivity of parameters. We use local and sobol^[22]^ method to calculate the sensitivity index of Teffs over-activation, Tregs activation and width of therapeutic window three indexes. (Fig. 5B shows and marked the parameters have significant impacts on the width of therapeutic window) These sensitivity indexes reflect the relative changings to target after a little parametric disturbance under this set of standard parameters. First-order sensitivity of parameters 41, 42, 30, 29 and total-effect index of parameters 14, 2, 17 are significantly higher than other parameters. Based on the treatment by rhIL-2, we may use other drugs to modify these specific parameters to enhance the efficacy of IL-2. The analysis of the sensitivity of parameters can pave the way for the establishment of an accurate medicine model in the future and can guide medication in clinic.

**Fig. 5.**
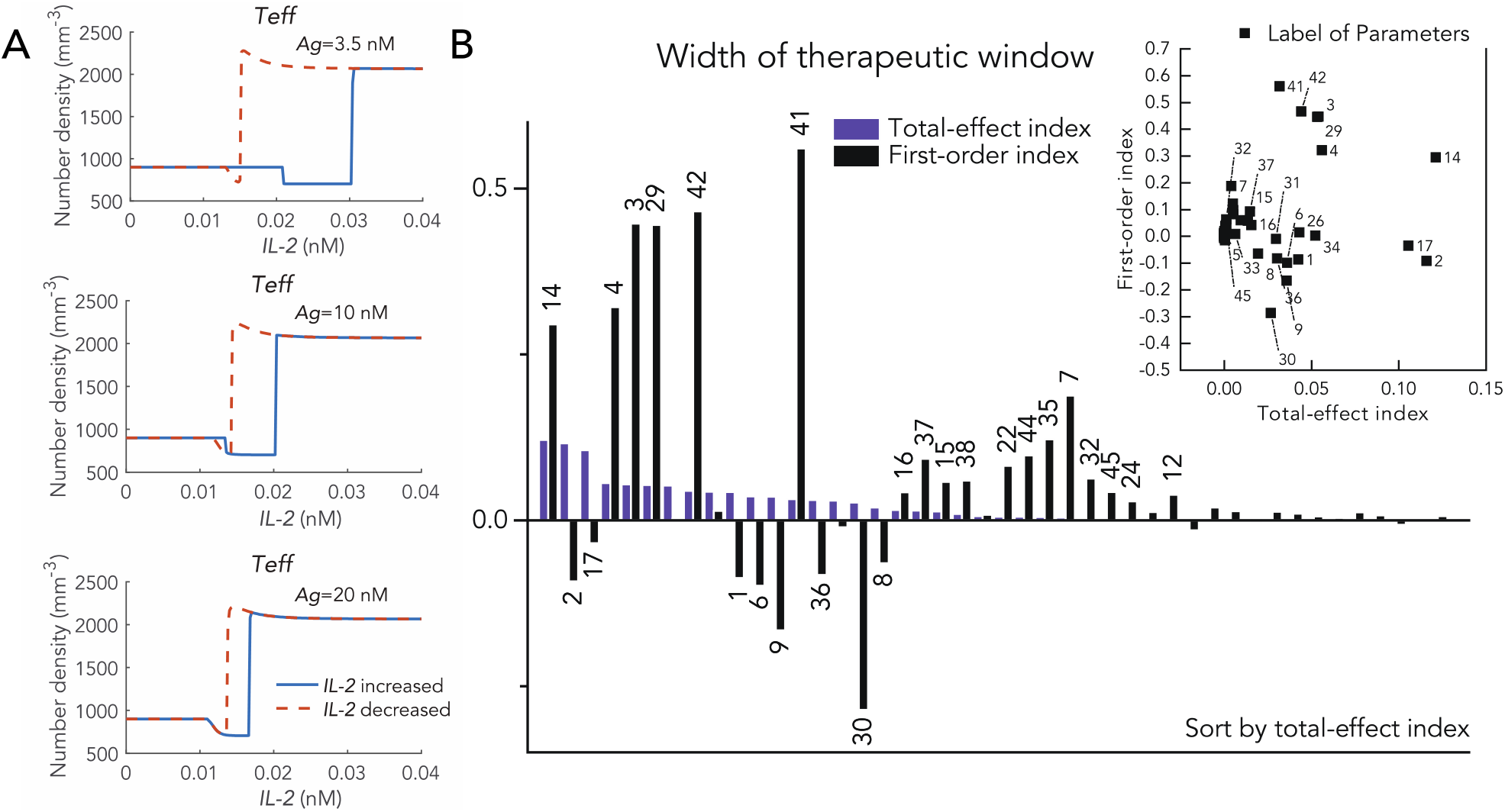
Parameters sensitivity analysis of therapeutic window. (A) Therapeutic window changes with different concentration of AgTCR (3.5 nM, 10 nM and 20 nM). (B) We exhibit the sensitivity of the width of therapeutic window by Variance-based sensitivity analysis (sobol method) and label the number of the corresponding parameters. The black column demonstrate the first - order sensitivity index and purple column represent the total-effect index. We sort the table by the absolute value of first-order sensitivity index and the descriptions of detailed parameters are list in table 1.

### Influence of environmental density on homeostasis: Lymph node and blood

Different organs of human have various immune tolerance to antigens, and the fact that lymph node is usually the first place to produce immune response when faced with virus or bacterial invasion reveal effect if the external environment on the immune response. Those organs that are rich in lymph nodes, such as the tonsils, will also have immune responses before than other organs. To illustrate the principle of this phenomenon, we modify other models after taking the volume of T cells into consideration and introduce a parameter β to represent the percentage of cell volume in the total volume(β=(V_*E*_ + V_*R*_)/V_*T*_). Of course, we also change the corresponding environmental load and constant multiplication rate to increase the cell density when reaches the steady state.

One of the distinction between lymph nodes and blood is the cell density and we simulate the proliferation and interaction of cells in different environments by changing cell density. We portray the concentration of endogenous IL-2 and soluble IL-2 with β and the ratio of Tregs and Teffs in Fig. 6A and B. It demonstrates that when the cell density increased gradually, the endogenous IL-2 concentration secreted by Teffs increase first with a sharp slope and then increasing slowly, but the soluble IL-2 concentration in body fluids tends to stabilize after a certain cell density (Fig. 6A). In addition, we also demonstrate that more intensive the cell density, the less antigens need to produce immune response (Fig. 6D-F) that reveals why the lymph nodes are the place the initial immune responses occurred. If we supply IL-2 in more intensive environment, however, more exogenous IL-2 needed to regulate the homeostasis compared to low-intensive environment like blood. Through the construction of the lymph node model, it can be convenient to simulate and predict the responses time of different organs in responses to antigens and finally facilitate clinical treatment.

**Fig. 6.**
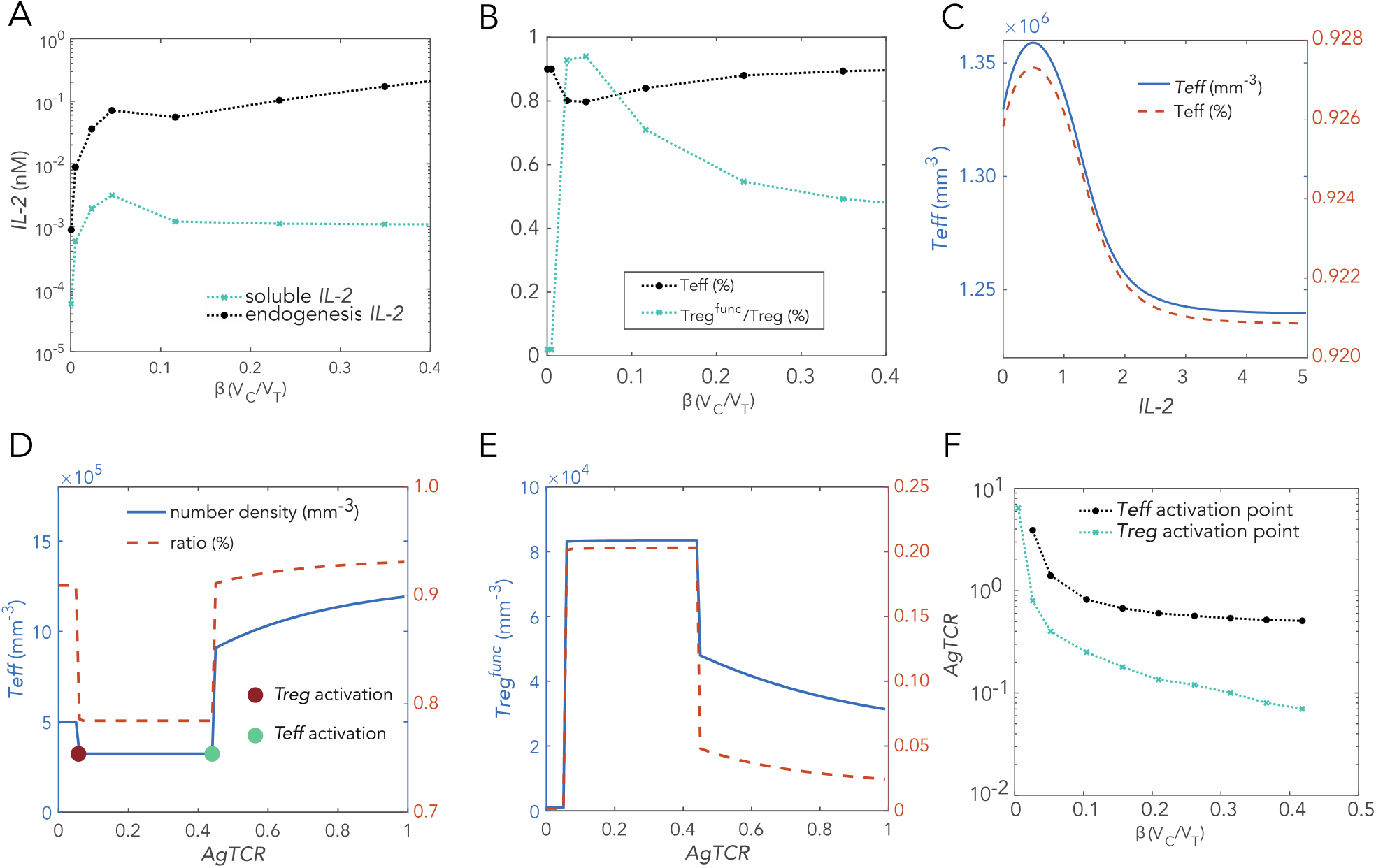
(A) The concentration of soluble IL-2 (blue dot line) and endogenous IL-2 (black dot line) vary with β (the proportion of total cell volume), which describes the differences between various kind of organs. (B) It describes the proportion of Teffs in CD4^+^ T cells and the proportion of functional Tregs in all Tregs with β. (C) When the initial cell density of Teffs was 1.33*106, the stable value of Teff will evolve with various concentration of exogenous IL-2. The left y-axis represents the change of Teff cell density, and the right y-axis represents the change in Teff fraction. (D-E) The Teff activation point (green dot) and Treg activation point (red dot) will appear in succession after stimulated by antigens. (F) It represents the concentration of antigens required for Teff and Treg activation in different reaction environments. The denser the reaction environment, the lower the concentration of antigen required for the immune response, which explains why lymph nodes response first in our immune system.

### Dynamic behaviors of high dose IL-2 and periodic administration

Many recent studies indicate the effectiveness of low dose IL-2 treatment in patients with dysfunctional Tregs. The kinetic behaviors of high does IL-2 treatment, however, are not clear yet, and the efficacy of the commonly used periodic administration in clinic has not been determined. Here, we report a simplified SLE model to investigate the dynamic behaviors of high dose and periodic administrations, which better characterizes the dynamic properties of the treatment process. Fig. 7A demonstrates the schematic diagram of simplified SLE model. Here, we identify a dynamical motif comprising the essential interactions of functional Treg, Teff and exogenous IL-2 and posit these as the central of SLE treatment. We developed the following mathematical model to analyze the treatment process:

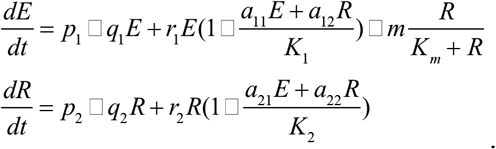

**Fig. 7.**
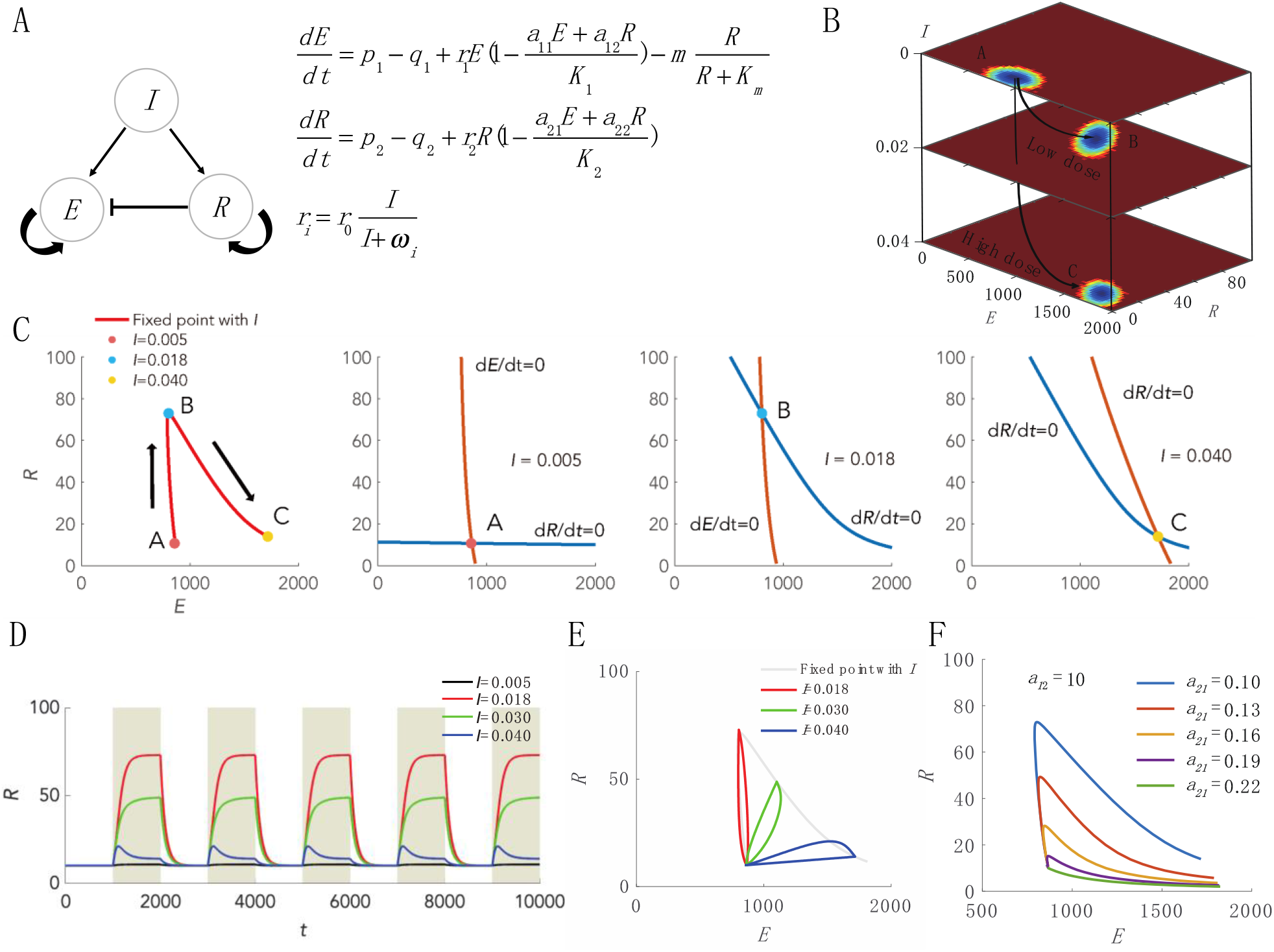
Dynamic behaviors of simplified SLE model and high dose IL-2 administration. (A) Schematic diagram of the simplified model and the ordinary differential equations. Symbol *I* represents the concentration of exogenous IL-2, *E* represents number density of Teff and *R* is the number density of Treg. (B) Dynamic landscape of the treatments with *I*. The horizontal coordinates denote the number density of *E* and *R*, the vertical coordinate represents the doses of exogenous *I* including 0, 0.02 and 0.04. Original, low dose and high dose attractors are labeled as A, B and C respectively. The black lines with arrow denote the paths from original attractor to low dose and high does attractors after treatment with exogenous *I*. (C) Phase diagrams at three doses of *I* and its nullclines. The left figure shows the stable fixed points with continuous doses of *I* from 0.005 (point A) to 0.040 (point C), and B denotes the dose of *I* at maximal value of *R* (about 0.018 in this set of parameters). The red lines denote the nullclines of *E* while blue lines denote the nullclines of *R*. (D) Kinetic evolution of *R* during periodic administration of four doses of *I*, and the brown shades denote the administration intervals. (E) The paths of periodic administrations plotted in fig. 7D on the phase diagram. (F) The changes of fixed point curve when taking different competing coefficients *a*_*21*_, which denotes the competitive effect of *E* on *R*. The fixed point curve, however, shows extreme sensitivity to *a*_*21*_..

We modeled the number density of Teff (*E*) and Treg (*R*), using a modified Lotka-Volterra (LV) equation^[23, 24]^ with a carrying capacity *K*_*i*_ (*K*_*1*_=2500, *K*_*2*_=200) and a proliferation rate *r*_*i*_ (*i*=1 denotes *E*, and *i*=2 denotes *R*), which depends on the dose of exogenous IL-2 *I* in the form of *r*_*i*_ = *r*_0_*I* / (*I* + *w*_*i*_) (*r*_*0*_=0.0389 denote the constant proliferation rate, and *w*_*i*_=[0.04, 0.01]denote the scaling constant). Teff cells are killed by functional Treg cells, *R*, using the Hill function with the maximal killing rate *m* and scaling constant *K*_*m*_=2. Matrix *a* in our model indicates the competition interactions between *E* and *R*, *a*_*11*_ and *a*_*22*_ indicate the effect of *E* and *R* on themselves (we set *a*_*ii*_ equals 1, *a*_*12*_=10 and *a*_*21*_=0.1). Furthermore, the competition interactions of Treg to Teff was proved strong than Teff to Treg, and we assume *a*_*12*_=10 and *a*_*21*_=0.1 respectively here. Besides, our mathematical model consists the proliferation without IL-2 with a constant proliferation rate *p*_*i*_ (*p*_*1*_=9, *p*_*2*_=0.1) and a degradation rate *q*_*i*_ (*q*_*1*_=0.01, *q*_*2*_=0.01).

The stable fixed point of this motif varies with exogenous dose of IL-2 (fig. 7c), point A, B and C denotes the stable state when *I* equals to 0.005, 0.018 and 0.040 corresponding to ultra-low, low and high dose of IL-2 respectively. Point B actually represents the dose of *I* at maximal value of *R*, and the ‘therapeutic window’ in our whole model refers to doses near point B. The nullclines of *E* and *R* clearly demonstrate the dynamical process of our motif. When *I* is small, the nullcline of *R* is sensitive while the nullcline of *E* is not, and the nullcline of R moves up to cause the stable point to evolve from A to B. When *I* is large, the nullcline of *E* is sensitive while *R* is not, the nullcline of E moves down to push the stable point move from B to C. The specific sensitivity of both nullclines create the therapeutic window in our motif, and make the scope of the therapeutic window fluctuates around point B with other clinic indicators. High dose of *I*, however, immediately cause the stable point to evolve from point A to C, and doesn’t pass anywhere around B (fig. 7B), which means high dose administration shields the therapeutic window and does nothing positive to the treatment.

We than investigate the dynamical process of periodic administration which frequently used in clinic and various researches. In some diseases, periodic administration gives the system time to recover, and avoid the development of drug resistant.^[24]^. According to fig. 7D and E, our model suggests that high dose of periodic medication can only be of limited help to the disease, although periodic administration can turn the trajectory around before it reaches homeostasis C, its direction is skewed away from the treatment window at the beginning. Different set of Parameters, however, will also influence the curve of stable points shown in fig. 7C. Competition matrix *a* which represents the interaction intensity between *E* and *R* has various values during different patients. We simulate five set of *a* in fig. 7F, it can be seen that the therapeutic window is highly sensitive to *a*_*21*_, which indicates that patient’s specificity during treatment is extremely important.

## Discussion

We presented a model of IL-2-Teff-Treg system to illustrate the distinct response of immune system to antigen and IL-2, and found it could not only be well fit to experimental data but recapitulate the distinct response by different IL-2 dose. In our model, we ignored the DC cells and peripheral Treg cell induction, which are said to be important in low antigen tolerance induce. However, one group has evidences that the maturity of DC cells are related to the IL-2-IL-2R complex on DC surfaces, therefore, the contribution of DC cells on tolerance induction could due to the level of IL-2 and IL-2R, which is the core pathway in our model. Besides, since Treg induction is another issue in mathematical modeling, and the time scale of differentiation process may be much longer than cell propagation process, we included the Treg cell differentiation process in initial ratio of Teff to Treg. By considering these complex interactions, our IL-2-Teff-Treg model can be an indicator of the bifurcation process of immune system by antigen dose and IL-2 dose^[25]^. For example, through the model we revealed that the forward movement of the patient’s Teff activation point undermines immune tolerance. As we all know, activation of immune system often requires the stimulation of antigen and enough Teff cell density simultaneously. When the immune system is activated, Treg cells are always activated first, and it actually enters a sub-activated state, which is activated, but not so strong. Teffs can only be activated if the antigen concentration is high enough to reach another threshold. This feature of the IL-2-Teff-Treg system ensures its tolerance to low stimulating antigens, and the forward movement of the patient’s Teffs activation point undermines this tolerance.

